# The role of elevated branched chain amino acids in the potent effects of vertical sleeve gastrectomy to reduce weight and improve glucose regulation in mice

**DOI:** 10.1101/2020.06.01.128157

**Authors:** Nadejda Bozadjieva Kramer, Simon S. Evers, Jae Hoon Shin, Sierra Silverwood, Yibin Wang, Charles F. Burant, Darleen A. Sandoval, Randy J. Seeley

## Abstract

Obesity and type 2 diabetes mellitus (T2D) are growing epidemics resulting in increased morbidity and mortality. An emerging body of evidence has shown that elevated levels of branched-chain amino acids (BCAA) and their metabolites are strongly positively associated with obesity, insulin-resistance and T2D. Bariatric surgery is among the best treatments for weight loss and the alleviation of T2D. Additionally, clinical studies have reported that bariatric surgery decreases the circulating levels of BCAA. The objective of these studies was to test the hypothesis that reduced BCAA levels contribute to the metabolic improvements after VSG. We find that, as in humans, circulating BCAA levels are significantly lower in VSG rats and mice compared to Sham controls. In order to increase circulating BCAA levels, we tested mice with either increased dietary intake of BCAA or impaired BCAA catabolism by total body deletion of mitochondrial phosphatase 2C, Pp2cm, a key enzyme in the rate-limiting step in BCAA catabolism. Our results show that a decrease in circulating BCAA levels is not necessary for sustained body weight loss and improved glucose tolerance after VSG. While it is clear that circulating levels of BCAAs are excellent biomarkers for metabolic status, the current data do not support a causal role in determining metabolic regulation and the response to VSG.

## Introduction

Obesity is a growing epidemic that affects nearly a third of the population in the United States. Co-morbidities associated with obesity, such as Type 2 Diabetes (T2D), have estimated annual costs of $147 billion in medical care in the US. Numerous studies have focused on understanding the metabolomics profile associated with obesity and its comorbidities. Consequently, metabolic signatures including increased levels of circulating branched chain amino acids (BCAA) have consistently exhibited a strong correlation with obesity and insulin resistance (1-8).

Human and rodent data have shown that BCAA (leucine, isoleucine, and valine) are increased in both obese humans and rodents (1, 9). Elevated plasma levels of these three essential amino acids are associated with 5-fold increased risk for future development of T2D (10). Improving BCAA catabolism with a pharmaceutical approach effectively lowers circulating BCAA levels and attenuate insulin resistance in obese mouse models, creating novel avenues for potential therapy (9). The strong link between increased BCAA levels and insulin resistance have been demonstrated in humans with metabolic disorders and higher BMIs. However, even in studies where humans are matched for BMI, those with insulin resistance have higher circulating BCAAs (3, 4, 11). Non-targeted metabolic profile of hyperglycemic/T2D and normoglycemic patients similarly revealed a strong correlation between BCAA and BCAA metabolites with impaired fasting glucose and T2D (12). These findings consistently demonstrate that perturbations in BCAA homeostasis are associated with metabolic dysfunction. The mechanisms behind these observations have been posited in several recent reviews on this topic (13-16). Importantly, these data have led to the hypothesis that increased BCAA levels directly contribute to metabolic dysfunction in obese and diabetic patients.

Although invasive, weight loss surgeries such as Roux-en-Y gastric bypass (RYGB) and Vertical Sleeve Gastrectomy (VSG), have proven to be among the best treatments for weight loss and T2D (17). Recent studies also showed that bariatric surgery significantly reduces circulating BCAA levels post-surgery and BCAA levels remain lower at 2 years post operatively (18-20). These effective surgical interventions not only lead to decreased circulating BCAA levels, but also have been shown to improve BCAA catabolism, specifically in adipose tissue (21, 22). Clinical data has also suggested that weight loss surgeries reduce the levels of circulating BCAA more effectively than conventional weight loss interventions (21, 23). However, these studies are difficult to interpret, because the diet consumed by the control group and weight-loss surgery patients are likely not the same. In fact, bariatric surgery alters food preference in humans and rodents (24, 25). Overall, it remains unclear whether decreasing (or normalizing) circulating BCAA levels in obese humans is a direct contributor to the metabolic improvements after these surgical interventions. Moreover, it is important to identify whether a decrease in circulating BCAA levels post-surgery could be a predicative measure of sustained body weight loss and improved glucose homeostasis.

VSG produces important weight-independent effects on metabolism, including increased GLP-1 secretion, early-phase insulin secretion, improved glucose tolerance and increased hepatic insulin sensitivity in rodent models (25-27). To answer the question whether improved BCAA homeostasis plays a role in metabolic improvements after bariatric surgery, we used rodent models of VSG, which have a decrease in BCAA coupled with profound improvement in weight loss and metabolic profiles. We used both dietary and genetic manipulations to prevent the VSG-induced reductions in circulating BCAAs in the context of high-fat diet (HFD)-induced obesity. First, we show that circulating BCAA levels are indeed lower in VSG compared to Sham rats and mice as soon as two weeks after surgery. Subsequently, we supplemented HFD with increased levels of the three BCAAs (leucine, isoleucine and valine) while maintaining the same total protein content as control HFD. Finally, we used a mouse model of impaired BCAA catabolism, by total body deletion of mitochondrial phosphatase 2C (Pp2cm). Pp2cm is a key enzyme and an activator of the mitochondrial branched-chain α-ketoacid dehydrogenase (BCKD) responsible for the rate-limiting step in BCAA catabolism. Our results showed that a decrease in circulating BCAA is not necessary for sustained body weight loss and improved glucose tolerance after VSG.

## Results

### Vertical Sleeve Gastrectomy decreases circulating levels of BCAA

Clinical studies show that weight-loss surgeries reduce the levels of circulating BCAA (18-21, 28). We first determined whether the decrease in circulating BCAA observed in humans is also observed in rodent models of VSG. We generated a cohort of Long-Evans rats with *ad lib* access to 45% Tso’s high-fat diet with butter fat before and after undergoing Sham or VSG surgery. Rats that received VSG had a significant decrease in body weight from 2-16 days post-surgery compared to rats that received Sham surgery (Figure 1A). On day 14, the percent and total levels of fasting BCAA were decreased in VSG compared to Sham animals (Figure 1B, C). The decrease in plasma BCAA levels was not attributed to decrease in total amino acid levels, which were actually increased in the VSG rats (Figure 1D). In addition to a decrease in plasma levels of valine, leucine and isoleucine in VSG rats, individual amino acid analysis show that VSG rats have increased levels of glutamic acid (Figure 1E). Glutamic acid, or glutamate, is produced during the transamination reaction, the first step in BCAA catabolism. Finally, the alterations in amino acid levels seen between VSG and Sham rats were independent of circulating insulin levels which were not different between the groups at the time of amino acid analysis (Figure 1F).

**Figure 1.**
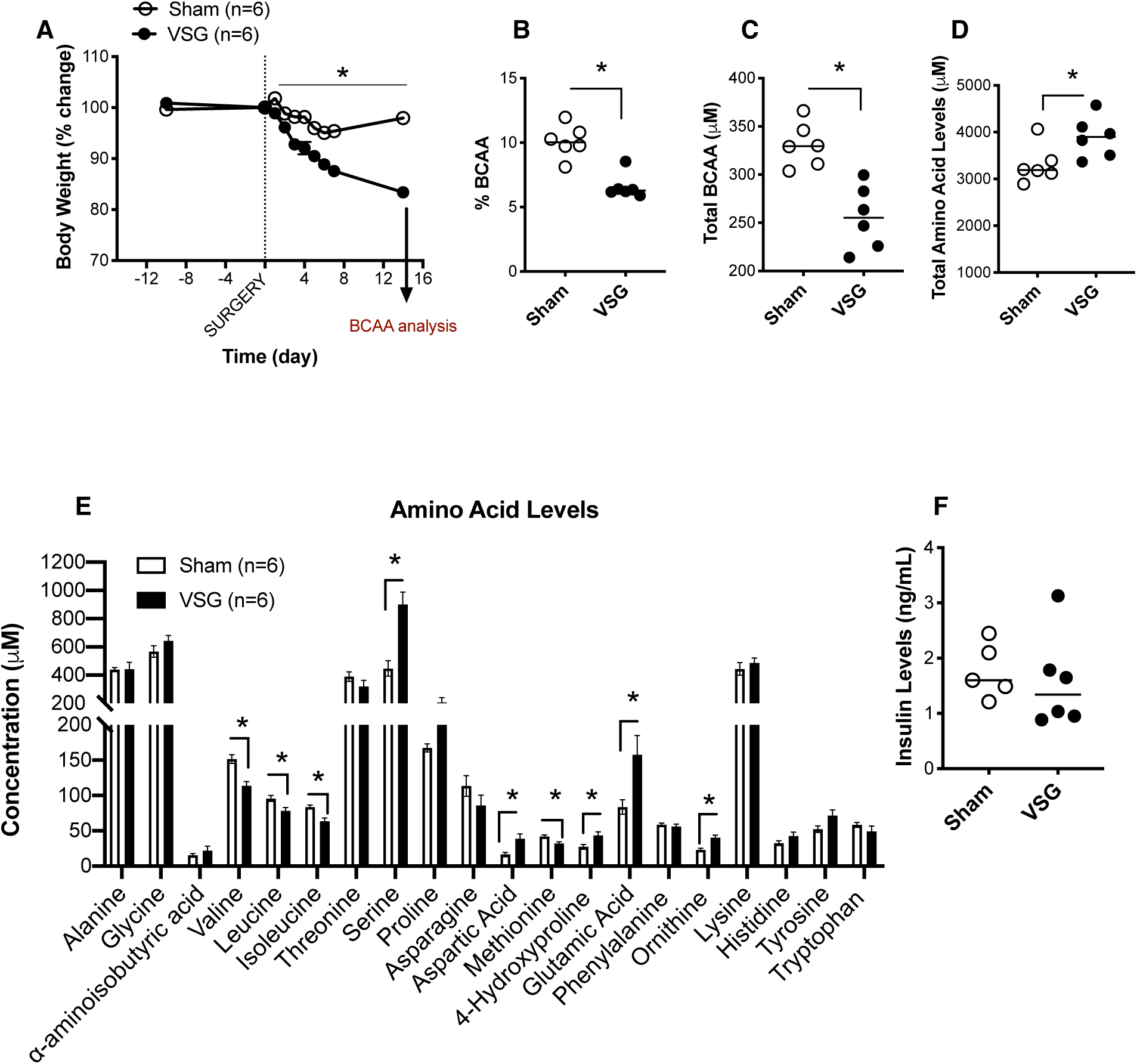
Vertical Sleeve Gastrectomy decreases circulating levels of BCAA. **A)** Body weight of Long-Evans rats with *ad lib* access to 45% Tso’s high-fat diet with butter fat before and after undergoing Sham or VSG surgery. **B)** Percent and **C)** Total levels of fasting plasma branched chain amino acids (overnight fast) on day 14 post-surgery. **D)** Total plasma amino acid levels on day 14 post-surgery. **E)** Individual plasma amino acid levels on day 14 post-surgery. **F)** Fasting insulin levels (overnight fast) on day 14 post-surgery. Animals n=6/group. Data are shown as means ± S.E.M. *p<0.05; (Student’s 2-tailed *t* test).

### Dietary supplementation of BCAA impairs glucose tolerance in high-fat diet (HFD) fed mice

We tested the impact of increased dietary intake of BCAA on body weight, adiposity, food intake and glucose tolerance in high-fat diet (HFD) fed mice. Wild-type male C57BL/6J were fed control and BCAA-supplemented HFD (Figure 2A). BCAA-supplemented diet contained 4 times the levels of BCAA compared to control diet (Figure 2A; Supplemental Table 1). The control and BCAA supplemented diet both contained 60% fat and had the same total amino acid content (Supplemental Table 1). Neither the control HFD nor the HFD+BCAA diet were deficient in any amino acids and essential fatty acids (Supplemental Table 2) (29). We included a small group of standard chow-fed mice as a reference group. Wild-type male C57BL/6J fed HFD and HFD+BCAA diets had comparable body weight gain, with similar adiposity and lean mass (Figure 2B, C, D). Mice fed HFD+BCAA showed decreased food intake 5 weeks after the start of the diet (Figure 2E). Additionally, they showed mild glucose intolerance after a mixed meal tolerance test (Figure 2F) and intraperitoneal tolerance test (ipGTT, Figure 2G) when compared to control mice fed HFD. BCAA supplementation did not affect fasting insulin or blood glucose levels (Figure 2H, I). The surrogate marker for insulin resistance, HOMA2 IR (calculated based on fasting blood glucose and insulin levels) was not different after dietary BCAA-supplementation during high-fat feeding compared to control HFD (Figure 2J).

**Figure 2.**
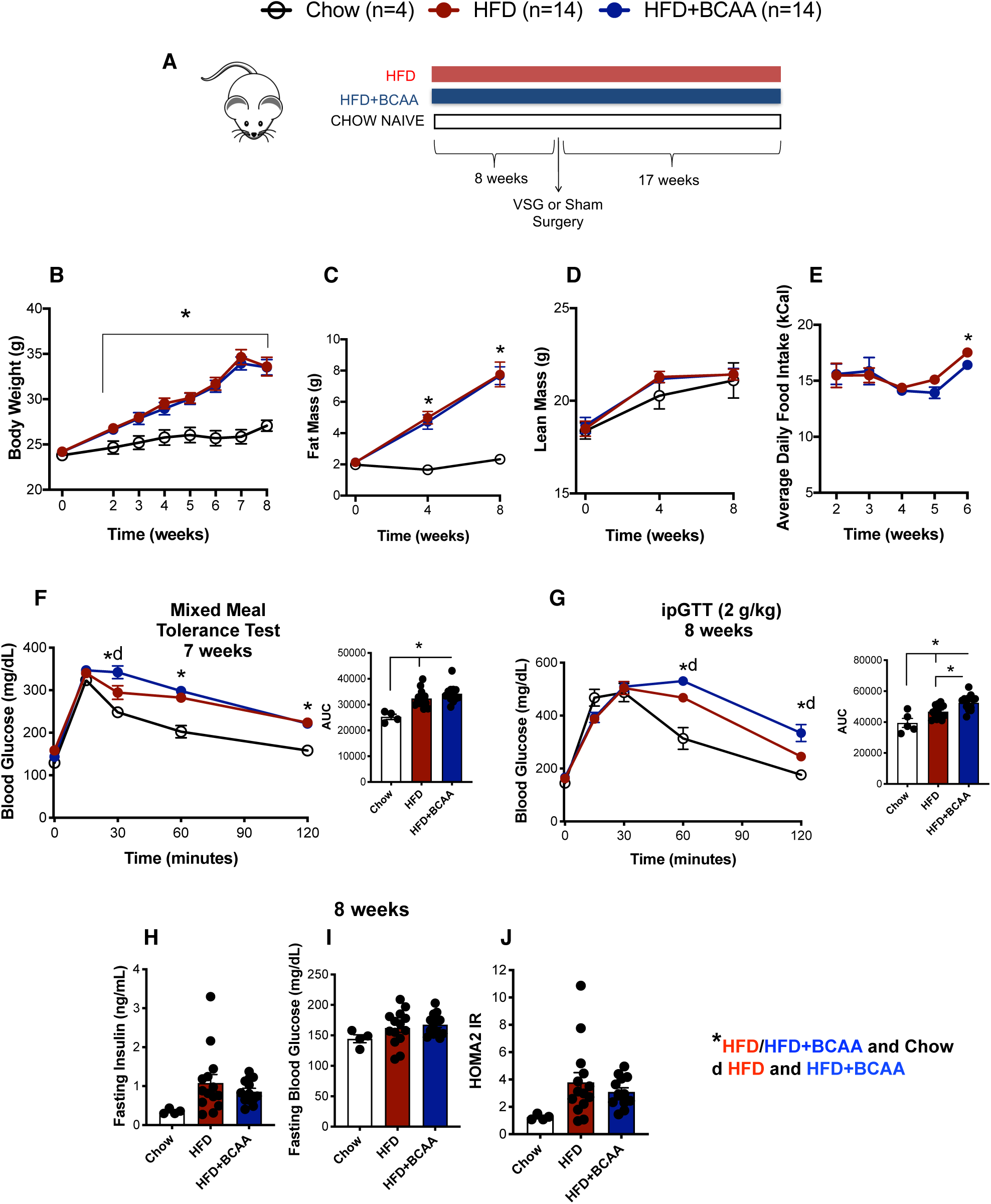
Dietary Supplementation of BCAA impairs glucose tolerance in high-fat diet (HFD) fed mice. A) Wild-type male C57BL/6J were fed control and BCAA supplemented high fat diet. BCAA-supplemented diet contained 4 times the levels of BCAA compared to the control diet. The control and BCAA supplemented diet both contained 60% fat and had the same total protein content (Supplemental Table 1). We included a small group of standard chow-fed mice as a reference group. **B)** Body weight, **C)** Fat mass, **D)** Lean mass, and **E)** Average daily food intake of male C57BL/6J fed HFD and HFD+BCAA diets. **F)** Mixed meal tolerance test was performed 7 weeks post initiation of diet. Shown as blood glucose response and area under the curve (AUC). **G)** Intraperitoneal tolerance test (ipGTT; 2 g/kg) was performed 8 weeks post initiation of diet. Shown as blood glucose response and area under the curve (AUC). **H)** Fasting insulin (6 hours), **I)** Fasting blood glucose levels (6 hours) and **J)** HOMA2 IR (calculated based on fasting blood glucose and insulin levels) 8 weeks after start of respective diet. Animals n=14/group. Data are shown as means ± S.E.M. * p<0.05 (1-Way ANOVA with Tukey’s post-test) with legends: **\*** p<0.05 HFD/HFD+BCAA compared to Chow; **d** p<0.05 HFD compared to HFD+BCAA.

### Dietary supplementation of BCAA does not impede sustained weight loss and improved glucose tolerance after Vertical Sleeve Gastrectomy (VSG)

Our results show that dietary BCAA supplementation along with high-fat feeding results in mild impairment in glucose tolerance. Next, we asked whether the decrease in BCAA levels after VSG contributed to the surgery-induced improvements in body weight loss and glucose tolerance. After 8 weeks of HFD or HFD+BCAA diet, mice were divided in four groups and underwent Sham or VSG surgery. Mice were returned to their pre-operative diet 3 days following Sham/VSG surgery. BCAA supplementation did not affect the sustained weight loss following VSG surgery (Figure 3A). We measured the levels of circulating amino acids after an overnight fast, 17 weeks after surgery. All four groups had comparable levels of fasting total plasma amino acid levels (Figure 3B). While the plasma BCAA levels (leucine, isoleucine and valine) showed only a trend towards being decreased in the VSG-HFD mice (Figure 3C), the percent of BCAA levels in plasma (relative to total amount of amino acid levels) was significantly lower in VSG-HFD mice compared to Sham-HFD (Figure 3D). This effect of VSG to lower BCAA levels was blocked in VSG-HFD+BCAA compared to Sham-HFD+BCAA (Figure 3C, D). BCAA supplementation did not affect the loss of fat mass following VSG surgery (Figure 3E). However, lean mass in VSG-HFD+BCAA was significantly lower compared to Sham-HFD+BCAA mice 13-16 weeks post-surgery (Figure 3F). Daily food intake decreased in both VSG groups after surgery. VSG-HFD had decreased daily food intake for 2 weeks after surgery, consistent to previous observations in wild type mice after VSG (30)(Figure 3G). VSG-HFD+BCAA mice had decreased daily food intake for 5 weeks post-surgery (Figure 3G).

**Figure 3.**
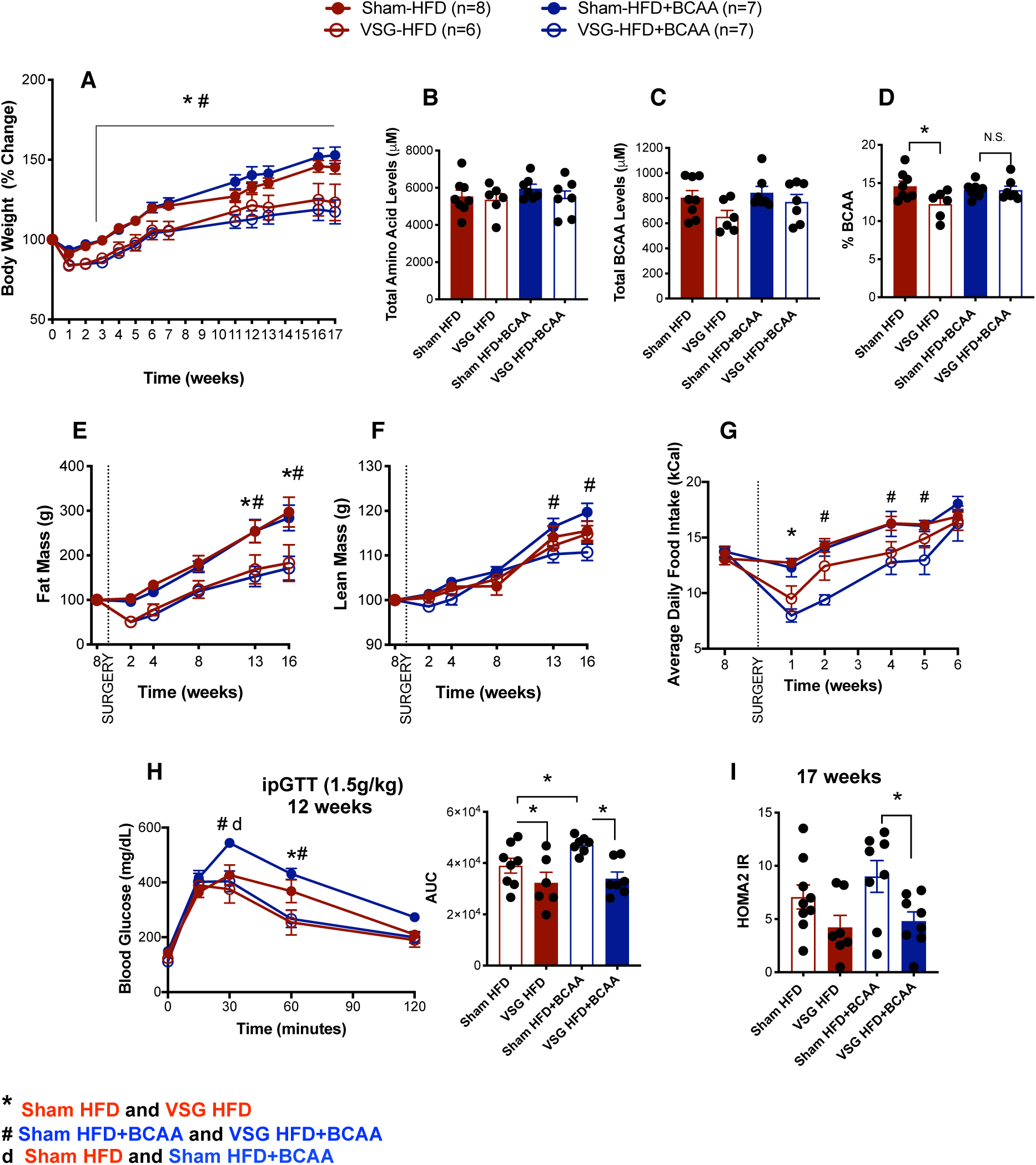
Dietary supplementation of BCAA does not impede sustained weight loss and improved glucose tolerance after Vertical Sleeve Gastrectomy (VSG). **A)** Body weight (% change) following VSG and Sham surgery. **B)** Total plasma amino acid levels, **C**) Total BCAA and **D**) Percent BCAA levels (relative to total amino acid levels) in plasma after overnight fast. **E**) Fat mass and **F**) Lean mass after VSG and Sham surgery. **G**) Average daily food intake after VSG and Sham surgery. **H**) Intraperitoneal tolerance test (ipGTT; 2 g/kg) was performed 12 weeks after VSG and Sham surgery. Shown as blood glucose response and area under the curve (AUC). I) HOMA2 IR, calculated based on fasting blood glucose and insulin levels, 17 weeks after surgery. Animals n=6-8/group. Data are shown as means ± S.E.M. * p<0.05 (2-Way ANOVA with Tukey’s post-test) with legends: **\*** p<0.05 Sham HFD compared to VSG HFD; **d** p<0.05 Sham HFD compared to Sham HFD+BCAA; **#** p<0.05 Sham HFD+BCAA compared to VSG HFD+BCAA.

Sham mice fed HFD+BCAA (20-week duration of diet, 12 weeks post-surgery) showed significant glucose intolerance after intraperitoneal tolerance test (ipGTT; 2 g/kg) when compared to control mice fed HFD (Figure 3H). Importantly, both VSG-HFD and VSG-HFD+BCAA had a significant improvement in glucose clearance compared to their respective Sham controls (Figure 3H). The surrogate marker for insulin resistance, HOMA2 IR (calculated based on fasting blood glucose and insulin levels after 6-hour fast) shows that VSG decreased insulin resistance after VSG independent of dietary BCAA-supplementation (Figure 3I). We observed a trend of higher HOMA2 IR in Sham-HFD+BCAA mice compared to Sham-HFD controls.

Indirect gas calorimetry measurements showed that dietary supplementation with BCAA lowered Respiratory Exchange Ratio after VSG. VSG-HFD+BCAA mice had decreased RER during dark phase compared to VSG-HFD mice (Supplemental Figure 1A-C). A trend of increased energy expenditure was seen in both VSG groups, with no differences between the two diets (Supplemental Figure 1D). No differences were seen in locomotor activity, food or water intake between the surgical groups and diets (Supplemental Figure 1E, F, G). Additionally, dietary supplementation with BCAA did not alter pancreas weight, pancreatic beta cell mass, liver and epididymal white adipose tissue (eWAT) (Supplemental Figure 2A-D). A trend of decreased pancreatic beta cell mass was seen in both VSG groups, with no differences between the two diets (Supplemental Figure 2B).

### Orally administered valine and glucose are absorbed faster into circulation in VSG compared to Sham rats

Our data show that increasing dietary intake of BCAA does not impede the sustained body weight loss and improvements in glucose tolerance after VSG. Subsequently, we tested the hypothesis that VSG decreases the absorbance of BCAA levels in the gut, thus leading to lower plasma BCAA levels. This hypothesis was supported by increased RNA expression of neutral amino acid transporters in ileal mucosal scrapes in VSG compared to Sham mice (Figure 4A). System L (*Slc7a5*/LAT1 and *Slc7a8*/LAT2) transporters release basolaterally smaller intracellular neutral amino acids in exchange for larger extracellular neutral amino acids (BCAA) (31, 32). These data suggested that the gut could also play a role in BCAA uptake through expressional regulation of amino acid transporters after VSG.

**Figure 4.**
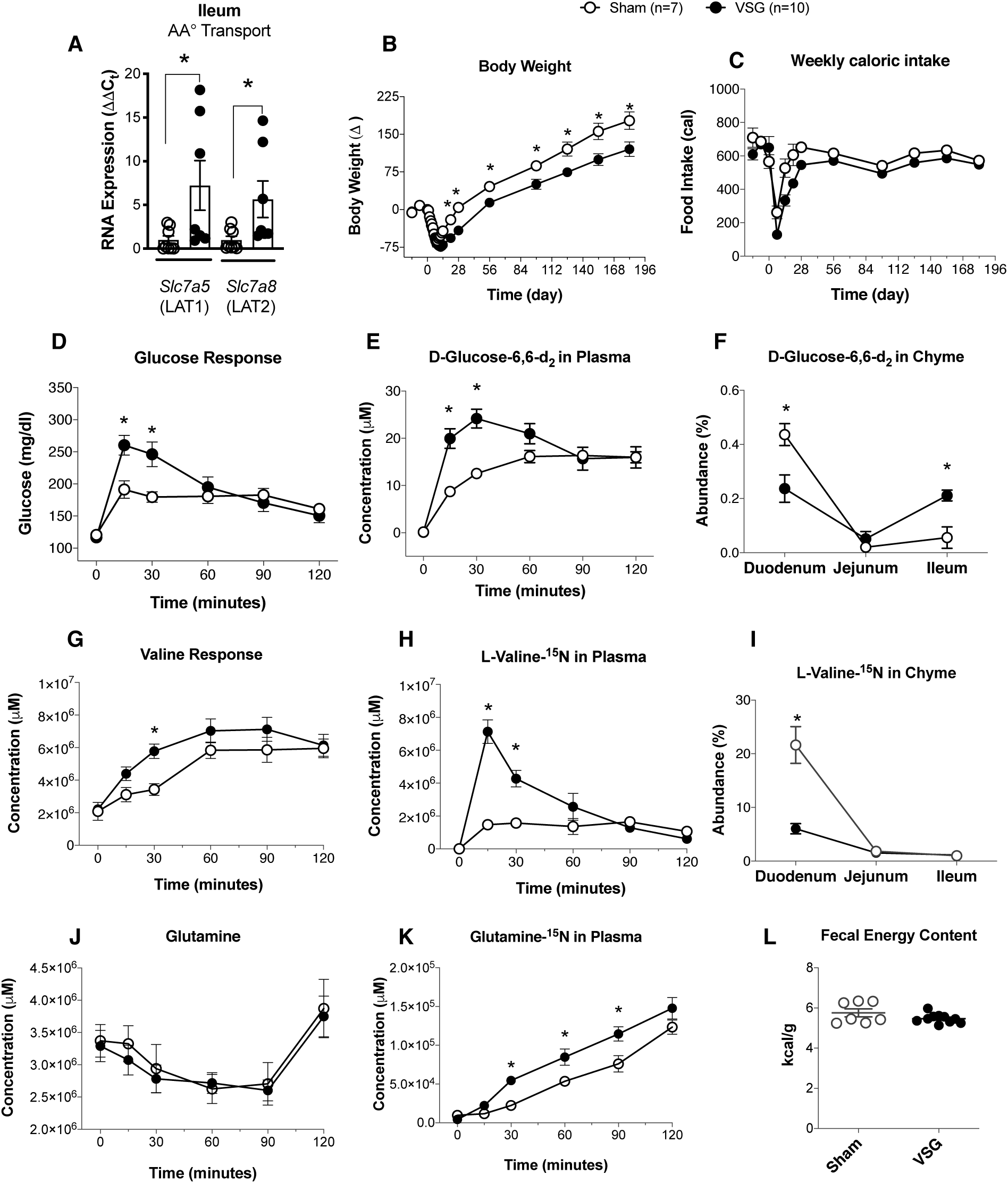
Orally administered valine and glucose are absorbed faster into circulation in VSG compared to Sham rats. **A)** RNA expression of amino acid transporters in ileal mucosal scrapes from wild-type Sham and VSG mice (n=6). **B)** Body weight and **C)** Weekly caloric intake in VSG and Sham Long Evans rats (n=7-10). **D)** Total glucose response, **E)** Circulating D-glucose-1,6-d_2_ and **F)** Luminal isotopic glucose in VSG and Sham Long Evans rats (n=7-10). **G)** Total circulating valine levels, **H)** Circulating levels of L-Valine-^15^N and **I)** Luminal levels of L-Valine-^15^N in VSG and Sham Long Evans rats (n=7-10). **J)** Total circulating glutamine levels, and **K)** Circulating levels of Glutamine-^15^N in VSG and Sham Long Evans rats (n=7-10). **L)** Fecal matter was collected from the period of 185-191 days post-surgery for energy quantification using bom-calorimetry (n=7-10) in VSG and Sham Long Evans rats (n=7-10). Data are shown as means ± S.E.M. * p<0.05; (Student’s 2-tailed *t* test).

To test the hypothesis that VSG leads to decreased absorbance of BCAAs into circulation, we generated a cohort of VSG and Sham Long Evans rats. Body weight and weekly caloric intake in VSG and Sham Long Evans rats show sustained body weight loss for 196 days following surgery, without differences in caloric intake (Figure 4B, C). At day 196, animals were fasted overnight and on the following morning received a mixture of liquid meal (Ensure Plus) containing 125mg/ml glucose + 125mg/ml 6,6-[^2^H]D-glucose + 12.5mg/ml [^15^N]L-Valine. VSG rats had a higher glucose and isotope-glucose excursion at 15 and 30 minutes following liquid meal, consistent with increased absorbance of glucose due to increased gastric emptying as a result of VSG (27, 33) (Figure 4D, E) and increased absorption capacity due to upregulation of the transporters *Slc7a5* and *Slc7a8* (Figure 4A). Labeled glucose levels were lower in duodenum chyme, but higher in the ileum chyme of VSG rats (Figure 4F). Similarly, VSG rats had a higher valine and isotope-valine excursion 15 and 30 minutes following liquid meal (Figure 4G, H). Labeled valine levels were lower in duodenum chyme of VSG rats compared to Sham rats (Figure 4I), consistent with increased efficiency of absorption. Glutamine levels were not different between Sham and VSG rats before (overnight fast) and following liquid meal (Ensure Plus) containing 125mg/ml glucose + 125mg/ml 6,6-[^2^H]D-glucose + 12.5mg/ml [^15^N]L-Valine (Figure 4J). However, labeled glutamine-^15^N, produced during the transamination reaction of valine, was increased in plasma of VSG rats compared to Sham (Figure 4K). Finally, fecal energy content was similar between Sham and VSG rats, demonstrating no malabsorption in VSG compared to Sham rats (Figure 4L). Overall, the data suggest that VSG leads to increased rate of valine absorption due to increased gastric emptying and increased rate of transamination.

### BCAA catabolism genes and amino acid transporter LAT1 expression is increased in epididymal white adipose tissue (eWAT) of VSG mice

Our data show that labeled glutamine-^15^N was increased in plasma of VSG rats compared to Sham after receiving a mixture of liquid meal (Ensure Plus) containing 125mg/ml glucose + 125mg/ml 6,6-[^2^H]D-glucose + 12.5mg/ml [^15^N]L-Valine (Figure 4K). Glutamine-^15^N is produced during the transamination reaction of valine and increased plasma levels of glutamine-^15^N suggested that VSG mice have increased BCAA catabolism. Therefore, we examined the expression of BCAA catabolism genes *BCDKHA* (BCDKH) and *BCAT2* (BCATm) in intestinal segments (duodenum, jejunum and ileum), liver (liver does not express *BCAT2*), soleus skeletal muscle and epididymal white adipose tissue (eWAT) of wild-type VSG and Sham mice (Figure 5A-F). White adipose tissue was the only site of increased expression of *BCDKHA* (BCDKH) and *BCAT2* (BCATm) in VSG mice, suggesting that VSG increases BCAA catabolism specifically in the eWAT. Additionally, expression of the amino acid transporter *Slc7a5* (LAT1) showed a trend of increased expression in the eWAT of VSG mice compared to Sham (Figure 5G). These data suggest that white adipose tissue has increased uptake and catabolism of BCAA following VSG.

**Figure 5.**
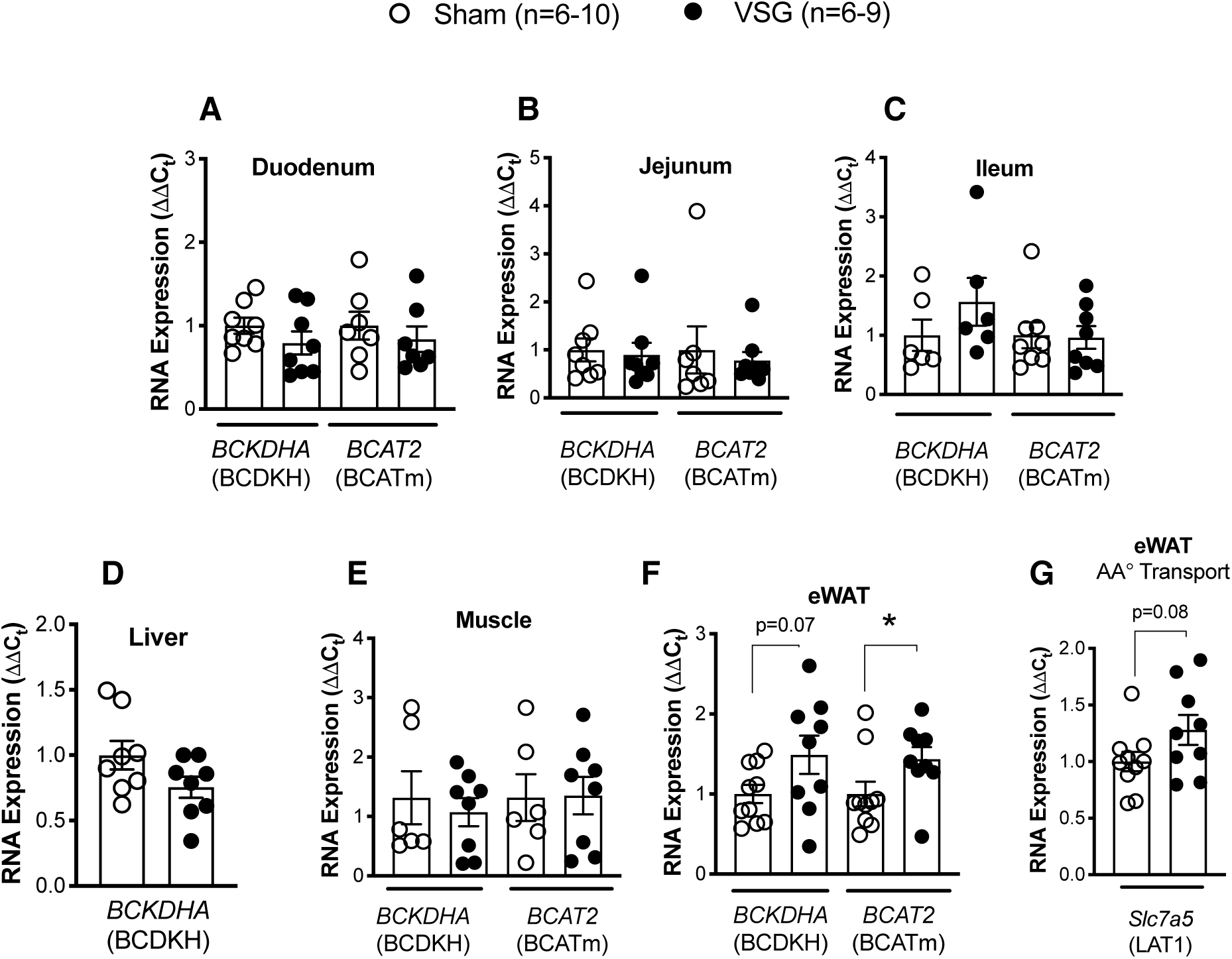
BCAA catabolism genes and amino acid transporter LAT1 expression is increased in epididymal white adipose tissue (eWAT) of VSG mice. RNA expression (ΔΔCt) of BCAA catabolism genes *BCDKHA* (BCDKH) and *BCAT2* (BCATm) in **A)** Duodenum, **B)** Jejunum, **C)** Ileum, **D)** Liver (liver does not express *BCAT2*), **E)** Muscle (soleus), **F)** Epididymal white adipose tissue (eWAT). **G)** RNA expression (ΔΔCt) of amino acid transporter *Slc7a5* (LAT1) in the eWAT of wild-type male Sham (n=6-10) and VSG mice (n=6-9). Data are shown as means ± S.E.M. **\*** p<0.05; (Student’s 2-tailed *t* test).

### Impaired BCAA catabolism, by total body ablation of Pp2cm, does not impede sustained body weight loss and improved glucose tolerance after VSG

Our data suggest that VSG mice have increased absorption, but also increased catabolism of BCAAs following VSG. Thus, we hypothesized that mice with decreased catabolism of BCAA will have impaired metabolic responses to VSG. To test this hypothesis, we utilized the Pp2cm^KO^ mouse model with a total body ablation of *Ppm1k* expression. *Ppm1k* encodes for mitochondrial phosphatase -- an activator of the mitochondrial branched-chain α-ketoacid dehydrogenase (BCKD) responsible for the rate-limiting step in BCAA catabolism (34). Consequently, genetically disrupted Ppm1k expression leads to partially impaired BCAA catabolism and elevated plasma levels of BCAA (34).

Pp2cm^KO^ mice at 6 weeks of age and under standard chow diet show a trend of lower lean mass, with no differences in body weight and fat mass (Supplemental Figure 3A-C). Intraperitoneal tolerance tests (ipGTT; 2 g/kg glucose) were performed in 6, 9, 12 and 30-week old chow fed Pp2cm^KO^ and WT mice and did not show differences in glucose clearance between the two genotypes (Supplemental Figure 3 D-G).

Pp2cm^KO^ and WT littermates gained comparable amount of body weight when challenged with 60% high-fat diet (HFD) for 8 weeks (Figure 6A). There were no genotype-specific differences in cumulative food intake over the 8-week span (Figure 6B).

**Figure 6.**
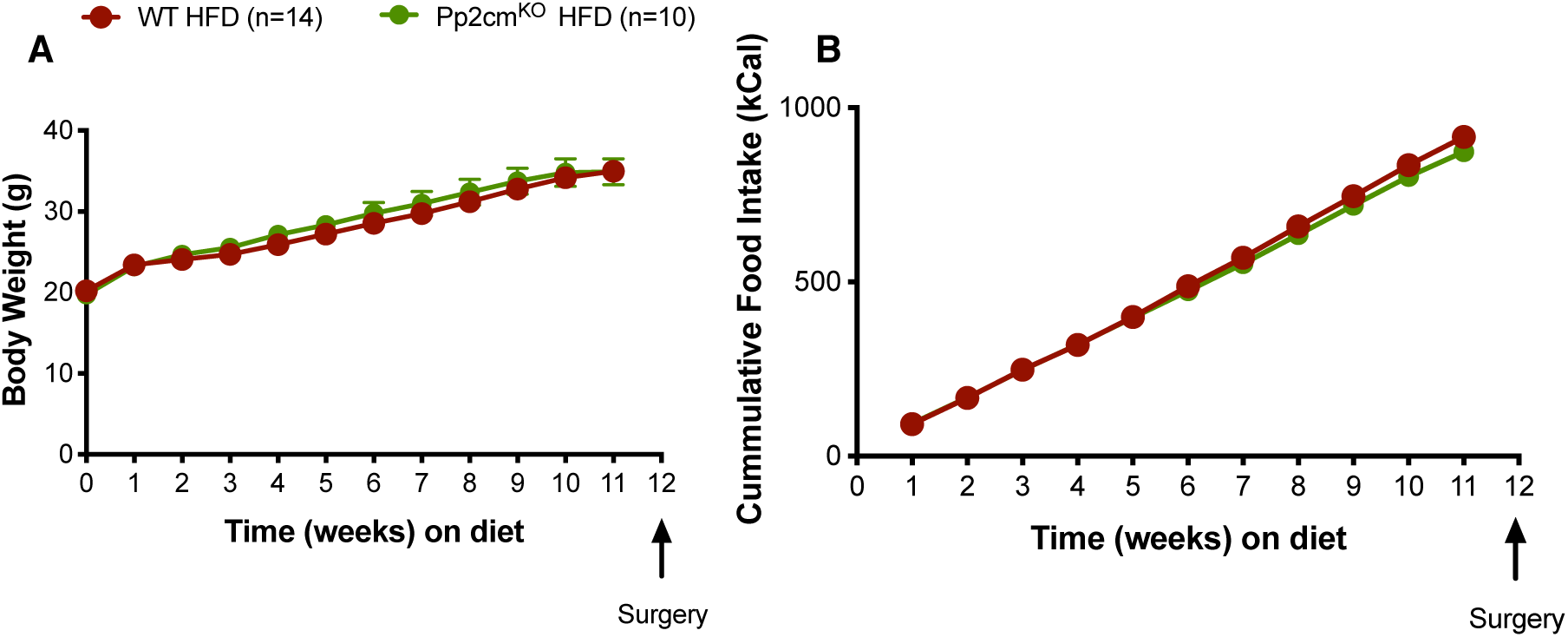
Impaired BCAA catabolism, by total body ablation of Pp2cm, does not lead to changes in body weight or food intake under standard chow or HFD conditions. **A)** Body weight and **B)** Cumulative Food intake in WT HFD (n=14) and Pp2cmKO HFD (n=10) mice fed 60% HFD for 12 weeks. Data are shown as means ± S.E.M. **\*** p<0.05; (Student’s 2-tailed *t* test).

In order to examine the effects of defective BCAA catabolism on body weight loss and glucose tolerance following VSG, we divided the mice into four groups that underwent Sham or VSG surgery. Mice were returned to their pre-operative diet 3 days following Sham/VSG surgery. Impaired BCAA catabolism, by Pp2cm ablation, did not affect the sustained weight loss following VSG surgery (Figure 7A). All four groups had comparable levels of total plasma amino acid levels (Figure 7B). Pp2cm^KO^ HFD-Sham mice had increased plasma levels of BCAA, and these levels did not decrease after VSG (Figure 7C). As expected, circulating BCAA levels were higher in Pp2cm^KO^ HFD-Sham mice compared to WT HFD-Sham mice (Figure 7C). Plasma BCAA levels and percent BCAA levels (relative to total amino acid levels) were also higher in Pp2cm^KO^ HFD-VSG mice compared to WT HFD-VSG mice (Figure 7C, D). Importantly, the ability of VSG to reduce BCAA levels was absent in Pp2cm^KO^ mice.

**Figure 7.**
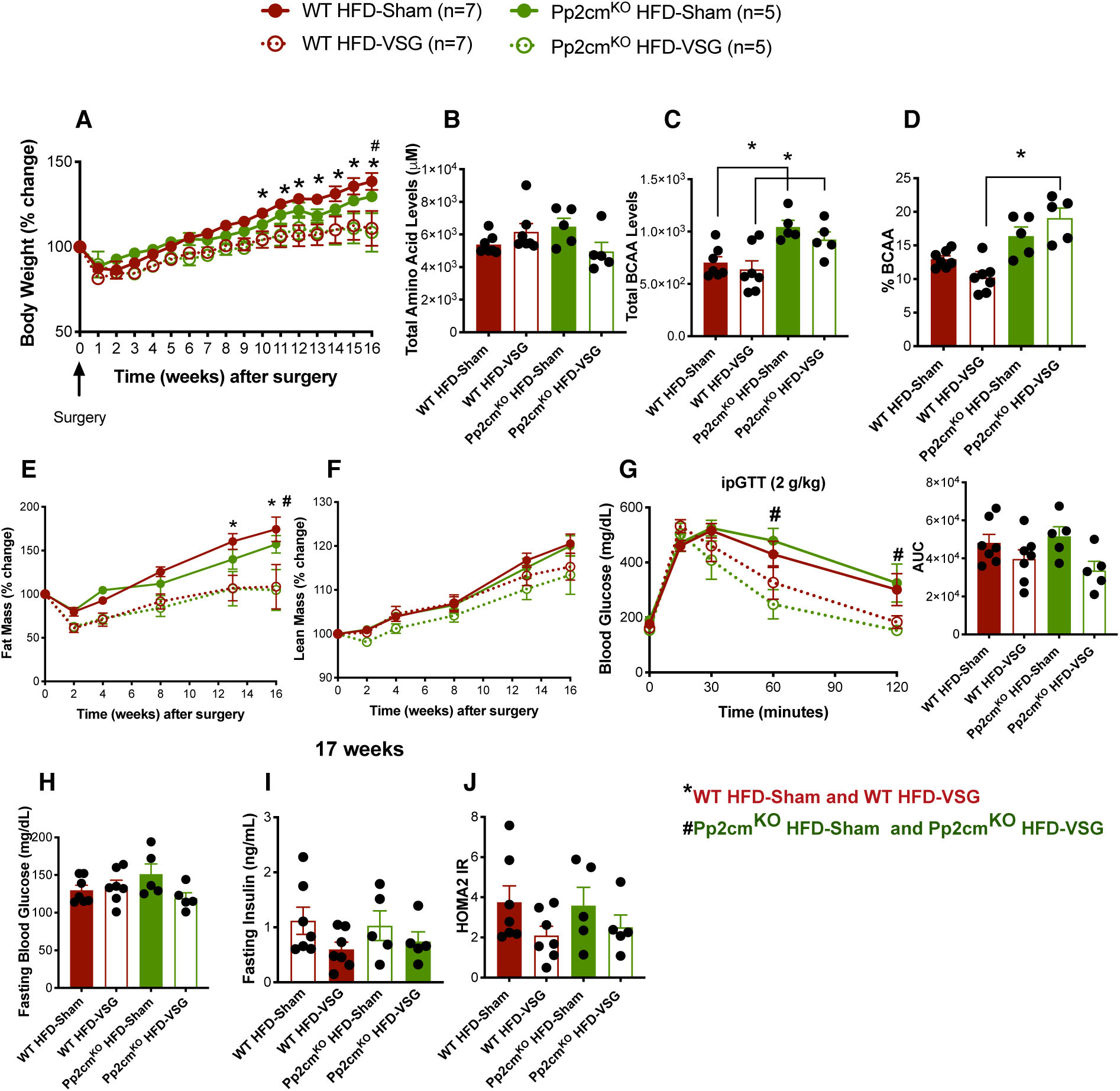
Impaired BCAA catabolism, by total body ablation of Pp2cm, does not impede sustained body weight loss and improved glucose tolerance after VSG. **A)** Body weight (% change) following VSG and Sham surgery. **B)** Total plasma amino acid levels, **C**) Total BCAA and **D**) Percent BCAA levels (relative to total amino acid levels) in plasma after overnight fast. **E)** Fat mass and **F)** Lean mass after VSG and Sham surgery. **G)** Intraperitoneal tolerance test (ipGTT; 2 g/kg) was performed 16 weeks after VSG and Sham surgery. Shown as blood glucose response and area under the curve (AUC). **H)** Fasting blood glucose levels (6 hours), **I)** Fasting insulin levels (6 hours) and **J)** HOMA2 IR, calculated based on fasting blood glucose and insulin levels, 17 weeks after surgery. Animals n=5-7/group. Data are shown as means ± S.E.M. * p<0.05 (2-Way ANOVA with Tukey’s post-test) with legends: * p<0.05 WT HFD-Sham compared to WT HFD-VSG; **#** p<0.05 Pp2cm^KO^ HFD-Sham compared to Pp2cm^KO^ HFD-VSG.

Defective BCAA catabolism did not affect the loss of fat mass following VSG surgery (Figure 7E). We observed a trend of decreased lean mass in Pp2cm^KO^ HFD-VSG mice, similar to what we saw in VSG-HFD+BCAA (Figure 3F and Figure 7F). Both VSG groups showed improved glucose clearance compared to their respective Sham controls, measured by intraperitoneal tolerance test (ipGTT; 2 g/kg) 16 weeks after surgery (Figure 7G). Fasting blood glucose levels were comparable between the four groups (Figure 7H). Although, not significant, fasting insulin levels were lower in both VSG groups compared to their respective Sham controls (Figure 7I). The surrogate marker for insulin resistance, HOMA2 IR (calculated based on fasting blood glucose and insulin levels after 6-hour fast), shows that VSG mice trended towards decreased insulin resistance after VSG independent of impairment in BCAA catabolism (Figure 7J). Pp2cm^KO^ mice HFD-Sham and HFD-VSG mice had similar meal size, meal number, food intake and locomotor activity compared to WT HFD-Sham and WT HFD-VSG mice (Supplemental Figure 4). These data suggest that impaired BCAA catabolism does not prevent the sustained body weight loss and improvement in glucose clearance following VSG.

## Discussion

Metabolomic profiling of obesity and insulin resistance compared to lean humans have identified branched-chain amino acids (BCAA) as a reliable biomarker of insulin resistance (1-8). Numerous clinical studies have shown that bariatric surgery reduces the circulating BCAA levels (18-21, 28). The objective of our study was to answer whether improved BCAA homeostasis is necessary for the metabolic improvements after VSG. First, we tested whether the decrease in circulating BCAA observed in humans is also observed in rodent models of VSG. Rats and mice that received VSG had a decrease in fasting BCAA levels as early as 2 weeks post-surgery (rat; Figure 1) and 17 weeks post-surgery (mouse; Figure 3). This decrease in plasma BCAA levels was not attributed to decrease in total amino acid levels. Also, at 2 weeks post-surgery, decreased levels of fasting plasma BCAAs was not due to reduced insulin levels, which were comparable between Sham and VSG rats.

Next, we tested the impact of increased dietary intake of BCAA on body weight, adiposity, food intake and glucose tolerance in high-fat diet (HFD) fed mice. Wild-type male C57BL/6J were fed control and BCAA supplemented high-fat diets, which contained four times the levels of BCAA compared to control diet (Supplemental Table 1). The control and BCAA supplemented diets both contained 60% fat and had the same total protein and carbohydrate content. Wild-type male C57BL/6J fed HFD and HFD+BCAA diets had comparable body weight gain, with similar adiposity and lean mass. Mice fed HFD+BCAA showed decreased food intake 5 weeks after the start of diet. Additionally, they showed mild glucose intolerance after a mixed meal tolerance test and intraperitoneal glucose tolerance test when compared to control mice fed HFD.

We sought to determine whether the decrease in BCAA levels after VSG contributed to the surgery-induced improvements in body weight loss or glucose tolerance. BCAA supplementation did not reduce the effectiveness of VSG to reduce food intake and produce sustained weight loss following VSG (Figure 3). The percent BCAA plasma levels (relative to total levels of amino acids) decreased in VSG-HFD mice compared to Sham-HFD. However, in VSG mice exposed to increased BCAA dietary supplementation, this decrease in circulating BCAA levels was greatly attenuated. The changes in plasma BCAA levels were not due to alterations in total plasma amino acid levels. All four groups had comparable levels of total plasma amino acid levels. Sham mice fed HFD+BCAA (20-week duration of diet, 12 weeks post-surgery) showed significant glucose intolerance when compared to control mice fed HFD (Figure 3). Importantly, both VSG-HFD and VSG-HFD+BCAA had a significant improvement in glucose clearance compared to their respective Sham controls (Figure 3). These data support the conclusion that dietary supplementation of BCAA does not impede sustained weight loss and improvement in glucose tolerance after VSG.

Indirect calorimetry measurements showed that dietary supplementation with BCAA lowers Respiratory Exchange Ratio (RER) after VSG. VSG-HFD+BCAA mice had decreased RER during dark phase compared to VSG-HFD mice (Supplemental Figure 1). These data suggest that VSG increases the preference for carbohydrates as metabolic substrates, but dietary supplementation with BCAA after VSG shifts the preference towards fats. Previous studies from our lab and others have shown that intracerebroventricular (ICV) infusion of leucine caused a decreased food intake in mice in an mTORC1-dependent manner (35-37). We did detect a decreased food intake in HFD+BCAA mice starting at around 5 weeks after BCAA diet initiation (Figure 2). Interestingly, after surgery the VSG-HFD+BCAA mice required more time to return their average daily food intake to levels comparable to their Sham-HFD+BCAA mice (Figure 3).

We did not observe increased fasting plasma BCAA levels in Sham-HFD+BCAA compared to Sham-HFD mice (Figure 3). Plasma BCAA levels spike, but decline back to baseline levels within 3 hours after consuming a protein-rich meal (38). We chose to measure the levels of BCAA under static conditions such as overnight fast, due to the differences in gastric volume and increased gastric emptying in VSG animals, which could have influence on the concentrations of plasma BCAA levels after feeding (27, 33). Consequently, we may have underestimated BCAA levels in the BCAA supplemented mice. Studies that have utilized diets with increased BCAA content have not reported plasma levels of BCAA after diet (39). Two recent studies also reported a lack of increase in the plasma BCAA levels after supplementing rodent diet with increased amount of BCAA and when compared to the standard diet (9, 40). This discrepancy makes it difficult to interpret if the impairment in glucose tolerance we and others see after dietary BCAA plus high-fat diet is due to 1) increased plasma levels of BCAA or 2) due to impaired BCAA catabolism resulting from high-fat feeding and increased BCAA diet load or 3) due to insulin resistance which may impair BCAA catabolism (41). Indeed, a recent Mendelian randomization study suggested that BCAA levels do not have a causal effect on insulin resistance in humans (42). Nevertheless, the current data support the hypothesis that the impairment in glucose tolerance in Sham-HFD+BCAA mice may be due in part to impaired catabolism of BCAA as a result of chronic exposure of increased BCAA content in the context of a HFD.

Increasing dietary intake of BCAA does not impede the sustained body weight loss and improvements in glucose tolerance after VSG. VSG does not decrease the absorbance of BCAA levels in the gut and we find no evidence of overall malabsorption given unchanged levels of fecal energy content. There is instead an of increased BCAA absorption, likely as a result of increased gastric emptying and increased RNA expression of neutral amino acid transporters in ileal mucosal scrapes in VSG compared to Sham mice (Figure 4). The L-type amino acid transporters (LAT) family, and in particular LAT1 (*Slc7a5*), is responsible for the transport and absorption of the majority of cellular BCAA (43). System L (*Slc7a5*/LAT1 and *Slc7a8*/LAT2) transporters basolaterally release smaller intracellular neutral amino acids in exchange for larger extracellular neutral amino acids (BCAA) (31, 32). Specifically, glutamine enters the cells in a Na+-dependent manner and imports the BCAA leucine into the cell via the Slc7a5-SLC3A2 transporter (44). These amino acid transporters (*Slc7a5*/LAT1, *Slc7a8*/LAT2) were increased in the ileal mucosa of VSG mice (Figure 4). To test this hypothesis, we administered a liquid meal containing 125mg/ml glucose + 125mg/ml D-glucose-6,6-d_2_ + 12.5mg/ml L-Valine-^15^N to Sham and VSG rats. VSG rats had a higher glucose and valine uptake, as well as higher isotope-glucose and isotope valine uptake, at 15 and 30 minutes following liquid meal. These data are consistent with increased gastric emptying rate observed after VSG (Figure 4) (27, 33). Potentially, the exposure of the small intestine to higher concentrations of BCAA due to increase gastric emptying may lead to the increased expression of the transporters, similar to that seen with increased dietary exposure to amino acids and other substrates (45).

Our results show that plasma BCAA levels are decreased in VSG rats and mice, and these changes are not due to decreased absorbance of BCAAs. Therefore, we assessed whether VSG improved BCAA catabolism in tissues, resulting in decreased plasma BCAA levels. Recent studies have shown that BCAA homeostasis is determined largely by the BCAA catabolic activities in tissues (46). BCAA catabolism begins with the initial transamination step, catalyzed by BCAA transaminase (BCAT) to produce branched chain keto acids. The second step of oxidative decarboxylation is catalyzed by branched chain α-ketoacid dehydrogenase (BCKD) complex, yielding CoA esters. BCKD complex is tightly regulated by the phosphorylation/inhibition by branched chain α-ketoacid dehydrogenase kinase (BCKDHK) and the dephosphorylation/activation by mitochondrial phosphatase 2C (Pp2cm).

Clinical data have shown that weight loss surgical interventions not only lead to decreased circulating BCAA levels, but also have shown to improve BCAA catabolism specifically in adipose tissue (21, 22). In agreement with these observations, VSG mice have increased expression of key catabolic genes (*BCDKHA* and *BCAT2)* in epididymal white adipose tissue. Clinical studies have supported these data, showing that patients who have undergone bariatric surgery have increased protein expression of BCATm in omental and subcutaneous fat (22). Additionally, expression of the amino acid transporter *Slc7a5* (LAT1) had a trend towards increased expression in the eWAT after VSG (Figure 5F). These data lead us to speculate that white adipose tissue has increased uptake and catabolism of BCAA following VSG.

Consistent with our hypothesis that VSG leads to increased BCAA catabolism, the expression levels of key catabolic genes *BCDKHA* and *BCAT2* were increased in eWAT in VSG mice compared to Sham controls. Also, our data show that labeled glutamine-^15^N, produced during the transamination reaction of valine, was increased in plasma of VSG rats compared to Sham after receiving the mixed meal with stable isotopes (Figure 4). Consequently, we tested whether mice with impaired catabolism of BCAA will have impaired metabolic improvements after VSG by whole-body knock out of mitochondrial phosphatase 2C (Pp2cm). The *Ppm1k* gene (coding for Pp2cm) was recently identified as a genomic region strongly associated with BCAA levels and T2D (47). Lack of Pp2cm only partially impairs BCAA catabolism but does result in increased levels of circulating BCAAs (34). Overexpression of *Ppm1k* (gene coding for Pp2cm) lowers circulating BCAA levels, reduces hepatic steatosis and improves glucose tolerance in the absence of weight loss in Zucker fatty rats (16, 48). We also chose to use a total body knock out model, instead of adipocyte-specific knock out, to prevent a compensatory shift of increased BCAA catabolism by other tissues. Finally, Pp2cm^KO^ have a normal lifespan and are not resistant to HFD-induced weight gain as other mouse models of impaired BCAA catabolism (49).

Pp2cm^KO^ and WT littermates gained comparable amount of body weight consuming comparable amount of food when challenged with 60% high-fat diet (HFD) for 8 weeks (Figure 6). After these mice were given either VSG or Sham surgery, Pp2cm^KO^ did not show the expected reduction in BCAAs typically seen after VSG (Figure 7). Despite the fact that BCAA’s remained high in Pp2cm^KO^ given a VSG, Pp2cm^KO^ showed the same sustained weight/fat loss following VSG surgery (Figure 7). Furthermore, Pp2cm^KO^ showed the same effect of VSG to improve glucose tolerance measured by an intraperitoneal tolerance test 16 weeks after surgery. These data show that impaired BCAA catabolism does not prevent weight loss and improvements in glucose caused by VSG.

BCAA supplementation or impaired BCAA catabolism by total body deletion of Pp2cm did not affect the loss of fat mass following VSG surgery. However, lean mass in VSG-HFD+BCAA was significantly lower compared to Sham-HFD+BCAA mice 13-16 weeks post-surgery (Figure 3). We also observed a trend toward decreased lean mass in Pp2cm^KO^ HFD-VSG mice compared to their Pp2cm^KO^ HFD-Sham controls (Figure 7F). Muscle loss is a common side-effect of rapid weight loss interventions, including bariatric surgery in humans (50-53). BCAA supplementation with high-fat diet induces a chronic activation of mTORC1 in skeletal muscle (1). Additionally, infusions of amino acids in humans activate mTORC1 and consequently decrease insulin sensitivity in muscle and liver (54, 55). A recent study used heavy isotope steady-state infusions to show that obese and insulin resistant *db/db* mice show a shift in tissue specific BCAA catabolism away from fat and liver and towards skeletal muscle (46). We speculate that chronic activation of mTORC1 (and the resultant inhibition of insulin receptor signaling) in the muscle of these mouse models (VSG-HFD+BCAA and Pp2cm^KO^ HFD-VSG) impairs the ability to maintain muscle mass in response to rapid weight loss caused by VSG.

Although, we observed decreased food intake and impaired glucose tolerance in mice fed HFD+BCAA as reported by others (1, 15, 35), we did not observe changes in markers of insulin sensitivity. The surrogate marker for insulin resistance, HOMA2 IR, shows that insulin resistance improves after VSG independent of dietary BCAA-supplementation or Pp2cm. However, we did observe a trend toward higher HOMA2 IR in Sham-HFD+BCAA mice compared to Sham-HFD controls (Figure 3I). It is possible that longer exposure to BCAA-supplemented HFD would have induced significant insulin resistance. Direct measurement of insulin sensitivity via hyperglycemic-euglycemic clamp would provide a more sensitive test of this hypothesis.

Chow-fed Pp2cm^KO^ mice showed a trend toward increased fat mass and lower lean mass, without differences in body weight (Supplemental Figure 3). Intraperitoneal tolerance tests in chow-fed Pp2cm^KO^ and WT mice did not show differences in glucose clearance between the two genotypes (Supplemental Figure 3). These findings differ from recently published studies by Wang and colleagues, showing that chow-fed Pp2cm^KO^ mice have improved glucose tolerance and insulin sensitivity (56). Wang and colleagues used wild-type C57BL/6 mice as controls and although the authors state that the controls are the same genetic background and aged-matched, it is unclear that these controls came from the same litters as Pp2cm^KO^ mice. The control mice used in our studies are male, wild-type littermates of Pp2cm^KO^ mice.

In summary, our results show that plasma BCAA levels are decreased in VSG rats and mice, and that these changes are not due to decreased absorption or improved total body catabolism of BCAAs. A decrease in circulating BCAA is not necessary for sustained body weight loss and improved glucose tolerance after VSG. The key question is whether there are common mechanisms that mediate both the alterations in BCAA regulation and improved glucose regulation that occur after bariatric surgical procedures. Answering how surgical manipulation of the gut can alter BCAA metabolism in non-gastrointestinal tissues is an important research goal that could lead to a better understanding of the gut’s involvement in BCAA regulation as well as identifying less invasive solutions to obesity and T2D.

## Experimental Methods

### Animals and Diet

Male mice were single-housed under a 12-hour light/dark cycle with ad libitum access to water and food. Eight-week-old wild-type (C57BL/6) mice were fed 60% HFD (D16121101; protein (18% kCal) carbohydrate (21% kCal) fat (61% kCal)) or 60% HFD supplemented with BCAA for 8 weeks prior to VSG or Sham surgery (D16121102; protein (18% kCal) carbohydrate (21% kCal) fat (61% kCal)) from Research Diets, New Brunswick, NJ. Diet formulations can be found in Supplemental Table 1. After surgery, the mice were returned to same experimental diets. Mice fed the standard chow diet were included as a reference (Envigo Teklad; catalog 7012). The diets were formulated and purchased from Research Diets, Inc. (New Jersey, US) and both diets are isocaloric (5.2 kCal/gm). HFD+BCAA have a 4-fold increase in BCAA (leucine, isoleucine and valine) but the same amount of total protein content as HFD control diet. Pp2cm^KO^ male mice (generous gift by Dr. Yibin Wang, UCLA) (34) were single-housed under a 12-hour light/dark cycle with ad libitum access to water and food. Pp2cm^KO^ mice were born in Mendelian ratios and were not distinguishable from their WT littermates. Eight-week-old Pp2cm^KO^ and WT male mice (all littermates) were fed 60% HFD from Research Diets, Inc. (New Jersey, US; Catalog D12492) for 8 weeks prior to VSG or Sham surgery and returned to HFD after surgery until end of study.

Long Evans male rats were maintained on 45% Tso’s HFD with butter fat (Catalog D03082706 from Research Diets, New Brunswick, NJ) prior to undergoing Sham or VSG surgery. Fecal matter was collected from the period of 185-191 days post-surgery for energy quantification using bomb-calorimetry (performed by University of Michigan Animal Phenotyping Core).

All animals were euthanized using CO_2._

### Indirect gas calorimetry and Body composition

Body composition was measured using an EchoMRI (Echo Medical Systems). Indirect gas calorimetry (animal’s oxygen consumption (VO2) and carbon dioxide production (VCO2) to estimate various metabolic parameters, including the respiratory exchange rate (RQ), energy expenditure, substrate utilization, food and liquid intake, and locomotor activity were measured using 24-cage TSE PhenoMaster system (TSE Systems; Germany).

### Vertical Sleeve Gastrectomy (VSG) in mice and rats

Mice were maintained on a 60% HFD and Long Evans male rats on a 45% Tso’s HFD for 8 weeks prior to undergoing Sham or VSG surgery, as described previously (33, 57, 58). See above for diet source and catalog numbers. Animals were anesthetized using isoflurane, and a small laparotomy incision was made in the abdominal wall. The lateral 80% of the stomach along the greater curvature was excised in VSG animals by using an ETS 35-mm staple gun (Ethicon Endo-Surgery). The Sham surgery was performed by the application of gentle pressure on the stomach with blunt forceps for 15 seconds. The day prior and 3 days following surgery, animals were placed on liquid diet (Osmolite 1.0 Cal, Abbott Nutrition). They were placed back on pre-operative solid diet on day 3 post-surgery. Body weight and food intake as well as overall health were monitored daily for the first 7 days after surgery and once weekly until end of the studies.

### Isotope Studies

Long Evans male rats (Envigo, Indianapolis, IN) were individually housed at arrival and had *ad lib* access to 45% Tso’s high-fat diet with butter fat (D03082706 Research Diets, New Brunswick, NJ) and water. Rats weighed 518±9.65 g at time of surgery: Sham (n=8) and VSG (n=12) as described previously. At day 196, food was removed at the start of dark phase (18:00hr, 12:12 light-dark cycle) and animals were fasted overnight. Animals received a mixture of Ensure Plus containing 125mg/ml glucose + 125mg/ml D-glucose-6,6-d_2_ (Sigma-Aldrich) + 12.5mg/ml L-Valine-^15^N (Sigma-Aldrich) at a volume of 8ml/kg and a dosage of 1g/kg glucose, 1g/kg D-glucose-1,6-d_2_, and 0.1g/kg L-Valine-^15^N. Circulating glucose levels were measured at 0, 15, 30, 60, 90, and 120 minutes post-gavage from tail blood using a conventional glucometer (Accu-Chek) and an additional 200µl blood sample was collected and centrifuged to collect ∼100ul plasma. Plasma samples were stored at −80°C until they were analyzed for circulating isotope levels. Animals were euthanized 120 minutes post-gavage using CO_2_. Intestines were collected and carefully separated into duodenum, jejunum, and ileum. Contents of each section were collected and stored at −80°C. All samples used for isotope analysis were sent to the metabolomics core at the University of Michigan for isotope analysis.

### Metabolic Studies

Body weight was monitored monthly for 8 weeks prior and 17 weeks after Sham/VSG surgery. Intraperitoneal glucose tolerance test (IPGTT) was performed by intraperitoneal (IP) injection of 50% dextrose (2g/kg) in 4-6 hour fasted male mice. Mixed-meal tolerance test (MMTT) was performed via an oral gavage of liquid meal (volume 200 µl Ensure Plus spiked with a 25-mg dextrose) in 4-6 hour fasted male mice. Blood was obtained from the tail vein and blood glucose was measured with Accu-Chek blood glucose meter (Accu-Chek Aviva Plus, Roche Diagnostics). HOMA2 IR was calculated based on fasting blood glucose and insulin levels (6 hours of fasting).

### Amino Acid and Insulin Measurements

Insulin levels (6-hour fasting) were measured with ELISAs (Crystal Chem) following the manufacturer’s instructions. At week 17 post VSG/Sham surgery in mice and 2 weeks post VSG/Sham surgery in rats (Figure 1), food was removed at the start of dark phase (18:00hr, 12:12 light-dark cycle), and mice were fasted overnight. Plasma amino acid level analysis was performed at Michigan Regional Comprehensive Metabolomics Resource Core (University of Michigan, Ann Arbor). Amino acid levels were analyzed in overnight fasted plasma using the Phenomenex EZfast kit. Samples are extracted, semi-purified, derivatized and measured by EI-GCMS using norvaline as an internal standard for normalization. All blood was collected via tail vein in EDTA-coated tubes.

### Quantitative Real-Time PCR

RNA was extracted from tissue samples using RNeasy isolation kit (Qiagen). Gene expression was performed by quantitative real time RT-PCR using Power SYBR Green PCR Mix (Applied Biosystems) using StepOnePlus detection system (Applied Biosystems) with a standard protocol including a melting curve. Relative abundance for each transcript was calculated by a standard curve of cycle thresholds and normalized to βActin. Primers were purchased from IDT Technologies.

### Statistical Analysis

The statistical analysis for comparisons between 2 groups was performed by unpaired (2-tailed) Student’s t test. One-way ANOVA with post hoc Tukey’s multiple comparisons test was used for comparisons among 3 groups. Two-way ANOVA with post hoc Tukey’s multiple comparisons test was used for comparisons among 4 groups. P values <0.05 were considered significant (GraphPad Prism 8.2.0).

### Study Approval

All protocols were approved by the University of Michigan (Ann Arbor, MI) Animal Care and Use Committees and were in accordance to NIH guidelines.

## Acknowledgements

The authors thank the surgeons for conducting mouse VSG (Alfor Lewis, Andriy Myronovych, Mouhamadoul Toure, and Diana Farris) and Kelli Rule, Jack Magrisso and Stace Kernodle for the technical assistance. We thank Michigan Regional Comprehensive Metabolomics Resource Core (DK097153), University of Michigan Animal Phenotyping Core (1U2CDK110678-01) and University of Michigan In-Vivo Animal Core (IVAC) (University of Michigan, Ann Arbor). This work was supported by 5T32DK108740 (NB), 5T32DK071212-12 (NB), DK107282 (DAS), HL140116 (YW), DK020572 (MDRC), DK089503 (MNORC).

## Author Contributions

NB and RJS conceived, designed, analyzed results and wrote the manuscript. NB, SE, JS, SS performed the experiments and analyzed results. SE, DAS and CFB, conceived, designed, and analyzed results from stable isotope experiments. YW generated mice. CFB helped with data interpretation and discussion. All authors edited manuscript. RJS provided final approval of the submitted manuscript.

