## Supplemental Tables and Figures for "The role of elevated branched chain amino acids in the potent effects of vertical sleeve gastrectomy to reduce weight and improve glucose regulation in mice"

**Supplemental Table 1.** Special diet composition formulated and purchased by Research Diets, Inc. (New Jersey, US).

| **Product** | **Control HFD (gm) D16121101** | **HFD+BCAA (gm) D16121102** |
|---|---|---|
| Casein, 80 Mesh | 100 | 100 |
| L-Cystine | 3.35 | 1.05 |
| L-Isoleucine | 4.2 | 16.8 |
| L-Leucine | 7.74 | 30.96 |
| L-Lysine | 6.5 | 2.04 |
| L-Methionine | 2.3 | 0.72 |
| L-Phenylalanine | 4.4 | 1.38 |
| L-Threonine | 3.35 | 1.05 |
| L-Tryptophan | 1.05 | 0.33 |
| L-Valine | 5 | 20 |
| L-Histidine | 2.3 | 0.72 |
| L-Alanine | 2.3 | 0.72 |
| L-Arginine | 3.15 | 1.0 |
| L-Aspartic Acid | 5.7 | 1.79 |
| L-Glutamic Acid | 18.4 | 5.78 |
| Glycine | 1.6 | 0.50 |
| L-Proline | 10.3 | 3.24 |
| L-Serine | 4.75 | 1.49 |
| L-Tyrosine | 4.65 | 1.46 |
| BCAA (including casein) | 16.94 | 67.76 |
| Non-BCAAs (including casein) | 74.10 | 23.27 |
| Total L-Amino Acids | 91.04 | 91.03 |
| Maltodextrin | 125 | 125 |
| Sucrose | 68.8 | 68.8 |
| Cellulose, BW200 | 50 | 50 |
| Corn Starch | 0 | 0 |
| Soybean Oil | 25 | 25 |
| Lard | 245 | 245 |
| Fiber | 50 | 50 |
| Mineral Mix, S10026 | 10 | 10 |
| DiCalcium Phosphate | 13 | 13 |
| Calcium Carbonate | 5.5 | 5.5 |
| Potassium Citrate, 1 H2O | 16.5 | 16.5 |
| Sodium Bicarbonate | 3.5 | 3.5 |
| Vitamin Mix, V10001 | 10 | 10 |
| Choline Bitartrate | 2 | 2 |
| FD&C Yellow Dye #5 | 0.05 | 0.025 |
| FD&C Red Dye #40 | 0 | 0.025 |
| Total grams of diet | 765.39 | 765.38 |
| Protein kCal% | 18 | 18 |
| Carbohydrate kCal% | 21 | 21 |
| Fat kCal% | 61 | 61 |

**Supplemental Table 2.** Special diet composition formulated and purchased by Research Diets, Inc. (New Jersey, US).

| **Amino Acid** | **Control HFD (gm/kg diet) D16121101** | **HFD+BCAA (gm/kg diet) D16121102** | **NRC Requirement (gm/kg diet)** |
|---|---|---|---|
| L-Cystine | 5.2 | 2.2 | |
| L-Isoleucine | 10.4 | 26.9 | 4 |
| L-Leucine | 20.5 | 50.8 | 7 |
| L-Lysine | 17.1 | 11.3 | 4 |
| L-Methionine | 6.3 | 4.3 | 5* |
| L-Phenylalanine | 11.3 | 7.3 | 7.6** |
| L-Threonine | 9.1 | 6.1 | 4 |
| L-Tryptophan | 2.8 | 1.8 | 1 |
| L-Valine | 12.6 | 32.2 | 5 |
| L-Histidine | 6.0 | 3.9 | 2 |
| L-Alanine | 6.3 | 4.3 | |
| L-Arginine | 8.0 | 5.2 | 3 |
| L-Aspartic Acid | 15.4 | 10.3 | |
| L-Glutamic Acid | 49.0 | 32.5 | |
| Glycine | 4.0 | 2.6 | |
| L-Proline | 25.1 | 15.8 | |
| L-Serine | 12.8 | 8.5 | |
| L-Tyrosine | 12.1 | 7.9 | |
| *Per NRC Guidelines, Cystine may replace 50-66.6% of the Methionine requirement | **Per NRC Guidelines, Tyrosine may replace 50% of the Phenylalanine requirement | Reference: Council, N.R. (1995). Nutrient Requirements of Laboratory Animals: Fourth Revised Edition. (Washington (DC)). | |

Supplemental Figure 1

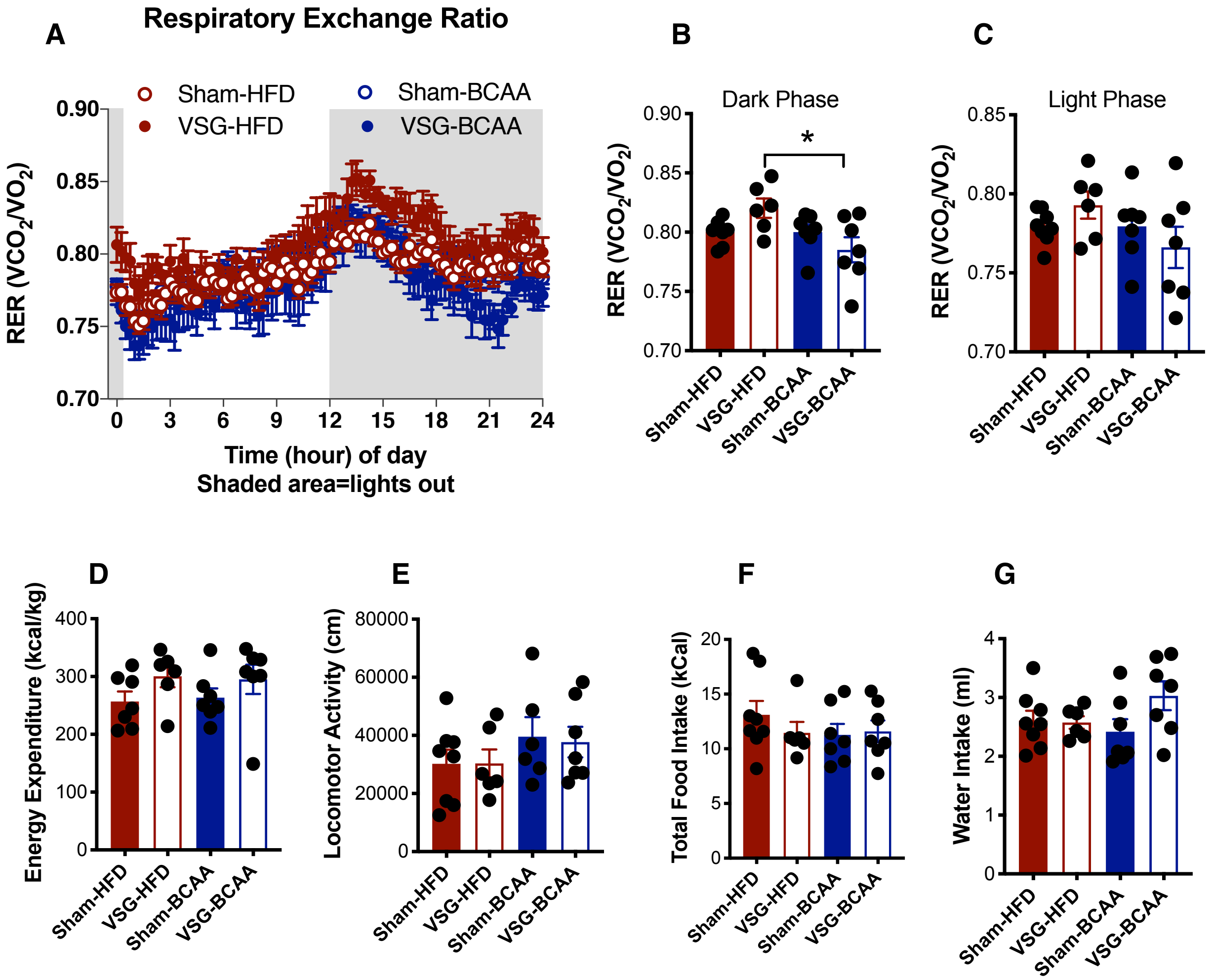

**Supplemental Figure 1. Dietary supplementation with BCAA lowers Respiratory Exchange Ratio after VSG. A)** Respiratory Exchange Ratio RER (VCO2/VO2) for 24-hour period. Respiratory Exchange Ratio RER (VCO2/VO2) for **B)** Dark and **C)** Light phase. **D)** Energy Expenditure (kCal/kg). **E)** Locomotor Activity (cm). **F)** Total Food intake (kCal) and **G)** Total Water Intake for Sham HFD (n=8), VSG-HFD (n=6), Sham-HFD+BCAA (n=7) and VSG-HFD+BCAA (n=7). Data represent average values for three consecutive days of recording. Shaded region in panel A represents dark phase. Data are shown as means ± S.E.M. * $p<0.05$; (2-Way ANOVA with Tukey's post-test).

Supplemental Figure 2

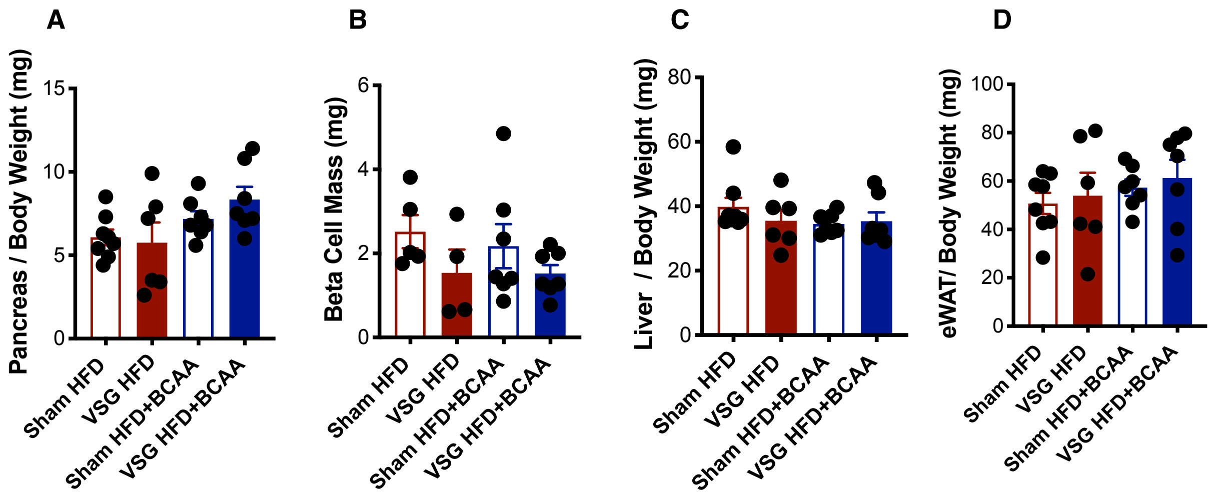

**Supplemental Figure 2. Dietary supplementation with BCAA does not alter pancreas weight, pancreatic beta cell mass, liver and epididymal white adipose tissue (eWAT). A)** Pancreas weights normalized to body weight in Sham HFD (n=8), VSG-HFD (n=6), Sham-HFD+BCAA (n=7) and VSG-HFD+BCAA (n=7). **B)** Pancreatic beta cell mass in Sham HFD (n=5), VSG-HFD (n=4), Sham-HFD+BCAA (n=7) and VSG-HFD+BCAA (n=7). **C)** Liver weights normalized to body weight in Sham HFD (n=8), VSG-HFD (n=6), Sham-HFD+BCAA (n=7) and VSG-HFD+BCAA (n=7). **D)** Epididymal white adipose tissue (eWAT) weight normalized to body weight in Sham HFD (n=8), VSG-HFD (n=6), Sham-HFD+BCAA (n=7) and VSG-HFD+BCAA (n=7). Data are shown as means ± S.E.M. * $p<0.05$; (2-Way ANOVA with Tukey's post-test).

Supplemental Figure 3

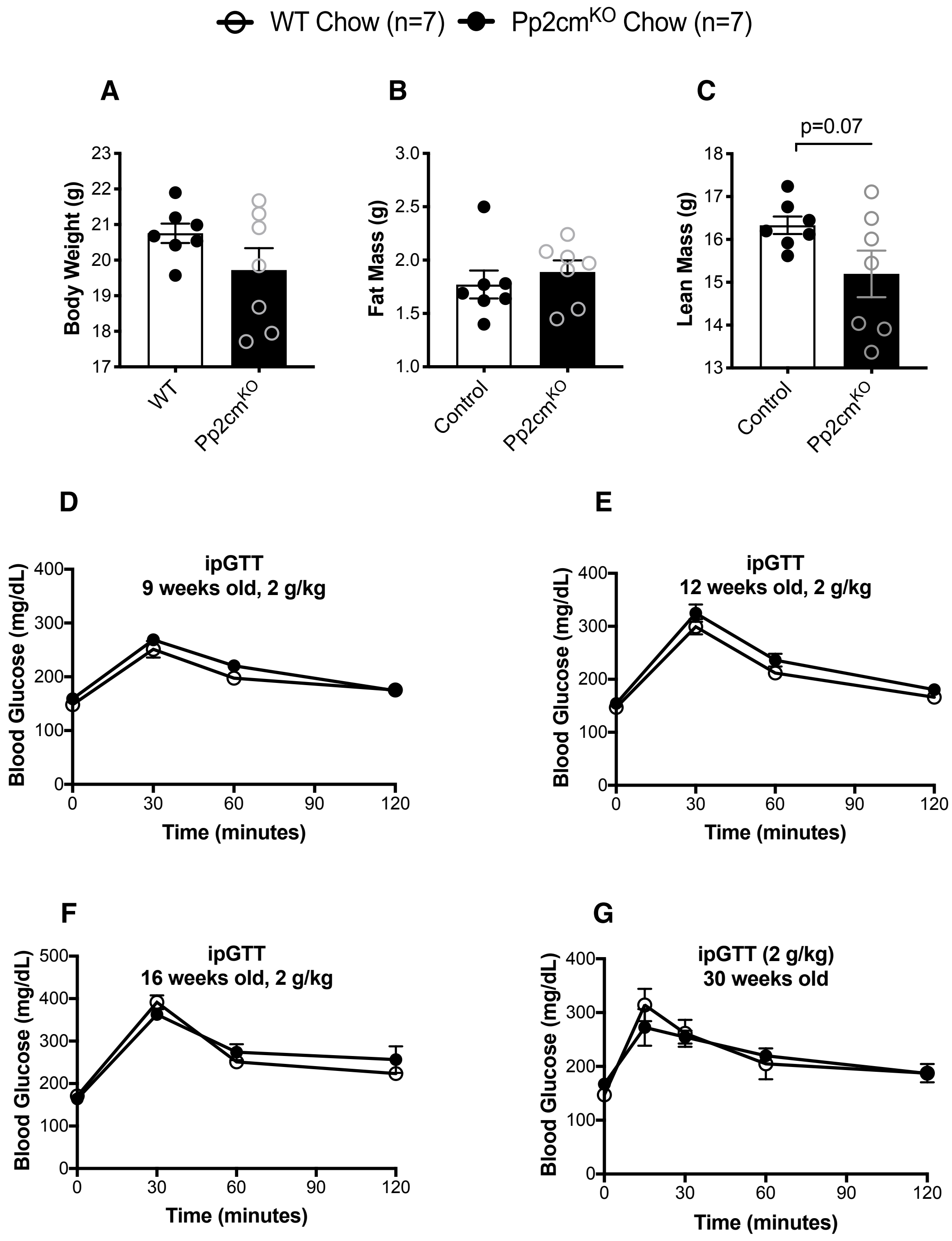

**Supplemental Figure 3. Impaired BCAA catabolism does not lead to glucose intolerance in standard chow-fed mice. A)** Body weight, **B)** Fat mass and **C)** Lean mass of 6-week old Pp2cmKO and littermate WT controls fed standard chow diet. Intraperitoneal tolerance tests (ipGTT; 2 g/kg) were performed in D) 9-weeks old, E) 12-weeks old, F) 16-weeks old and G) 30-weeks old standard chow-fed Pp2cmKO and littermate WT controls (n=7/genotype). Data are shown as means ± S.E.M. * $p<0.05$; (Student's 2-tailed *t* test).

Supplemental Figure 4

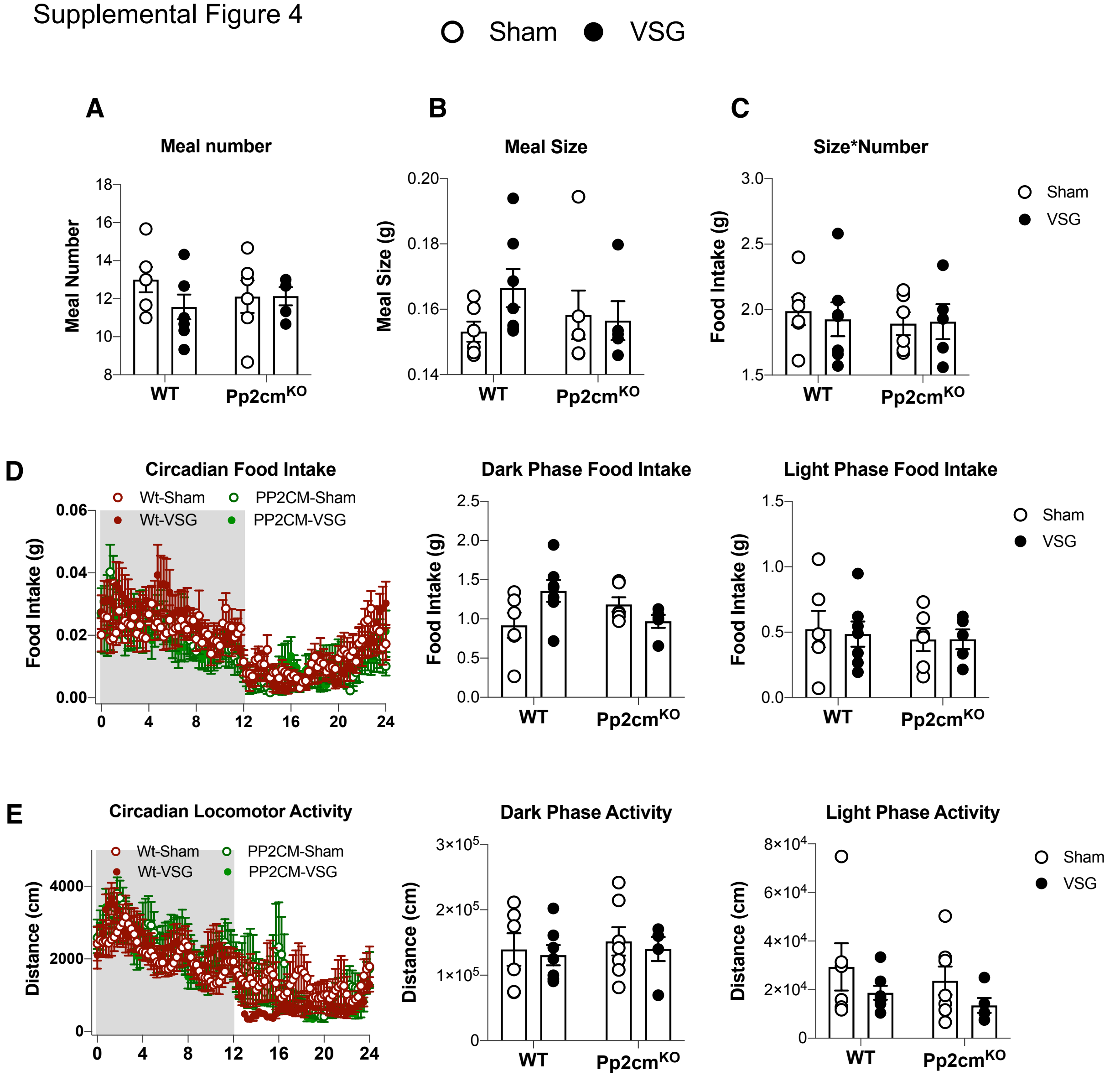

**Supplemental Figure 4. Impaired BCAA catabolism does not change food intake and locomotor activity in Sham and VSG mice. A)** Meal size, **B)** Meal number, and **C)** Meal size*Number. **D)** Circadian, **E)** Dark phase, and **F)** Light phase food intake. **G)** Circadian, **H)** Dark phase, and **I)** Light phase locomotor activity. Animals: WT HFD-Sham (n=7), WT HFD-VSG (n=6), $Pp2cm^{KO}$ HFD-Sham (n=5), $Pp2cm^{KO}$ HFD-VSG (n=5). Data are shown as means ± S.E.M. * $p<0.05$; (2-Way ANOVA with Tukey's post-test).
